# UDP-glucose pyrophosphorylase 2, a regulator of glycosylation and glycogen, is essential for pancreatic cancer growth

**DOI:** 10.1101/2020.10.13.337998

**Authors:** Andrew L. Wolfe, Qingwen Zhou, Eneda Toska, Jacqueline Galeas, Angel A. Ku, Richard P. Koche, Sourav Bandyopadhyay, Maurizio Scaltriti, Carlito B. Lebrilla, Frank McCormick, Sung Eun Kim

## Abstract

Pancreatic ductal adenocarcinomas (PDACs) have enhanced nutrient uptake requirements and rapid metabolic processing. The enzyme UDP-glucose pyrophosphorylase 2 (UGP2) rests at the convergence of multiple metabolic pathways, however the role of UGP2 in tumor maintenance and cancer metabolism remains unclear. Here, we identify an essential role for UGP2 in the maintenance of PDAC growth in both *in vitro* and *in vivo* tumor models. Transcription of UGP2 is directly regulated by the YAP/TEAD complex. Loss of UGP2 leads to decreased intracellular glycogen and defects in N-glycosylation targets important for cell growth including epidermal growth factor receptor (EGFR). In murine xenograft models, knockdown of UGP2 halted tumor growth and repressed expression of EGFR. The critical roles of UGP2 in cancer maintenance, metabolism, and protein glycosylation may offer new avenues of therapy for otherwise intractable PDACs.

**Impact Statement:** Convergent findings reveal that UDP-glucose pyrophosphorylase 2 has a central role in growth and metabolism of pancreatic ductal adenocarcinomas, highlighting novel therapeutic possibilities for this deadly cancer.

## Introduction

The majority of pancreatic ductal adenocarcinomas (PDACs) are mutant KRAS-driven tumors that are refractory to existing therapies, leading to a dismal 5-year survival rate below 10%^1^. As PDACs grow and expand their biomass, they must adapt to thrive in an increasingly dense, nutrient-limited, and hypoxic pancreatic environment. Tumors can adapt by rewiring their metabolism via stimulating scavenging pathways and upregulating rate-limiting enzymes involved in anabolic pathways^2,3^. For instance, the increase in glycolytic flux driven by this metabolic reprogramming can enhance cancer cell survival and drive chemoresistance^4,5^.

A key output of cellular metabolism is the manufacture of glycans, which are carbohydrate chains with fundamental roles in an array of cellular processes. Asparagine (N)-linked glycosylation changes can alter protein behavior, including deregulation of protein folding, enzymatic activity, and subcellular localization^6^. Recurrent changes in glycan modifications have been observed in cancer, although the precise patterns and regulation of these glycosylation changes remains unclear^7,8^.

The metabolism-regulating protein UDP-glucose pyrophosphorylase 2 (UGP2) is the only enzyme capable of converting glucose-1-phosphate to uridine diphosphate glucose (UDP-glucose) in mammalian cells. Its product UDP-glucose is upstream of both protein glycosylation and glycogen synthesis^9^. UGP2 is upregulated in some cancers, including PDACs, and its expression is correlated with an increased rate of progression and poor prognosis^10,11^. The direct role of UGP2 in growth and maintenance of PDAC has not been explored and the mechanisms of regulation and action of UGP2 in cancer remain unknown.

Here we determine that UGP2 is a key mediator required for the survival and proliferation of PDAC cells *in vitro* and *in vivo*. We identify that the transcriptional modifier Yes-associated protein 1 (YAP) is an important upstream regulator of UGP2 expression. We characterize two regulatory functions of UGP2 in PDAC cells: firstly, that UGP2 is critical for an array of protein N-glycosylation post-translational modifications including key sites on the epidermal growth factor receptor (EGFR), and secondly, that UGP2 regulates cellular glycogen synthesis under nutrient-starved conditions. These findings reveal metabolic pathways key for pancreatic cancer and provide potential clinical targets for further investigation.

## Results

### UGP2 is critical for PDAC cell survival and proliferation *in vitro* and *in vivo*

High expression of UGP2 correlated with worse clinical prognosis in a panel of 177 PDAC patient samples (Fig. S1A). To ascertain whether UGP2 is functionally required for the survival and proliferation of PDAC cells, UGP2 expression was depleted in the KRAS mutant pancreatic cancer cell line Suit2 along with several other pancreatic cancer cell lines (Fig. 1A-B and Fig. S1B-G). In each system, knockdown of UGP2 led to significant growth inhibition in both two-dimensional (2D) and three-dimensional (3D) culture settings.

**Figure 1.**
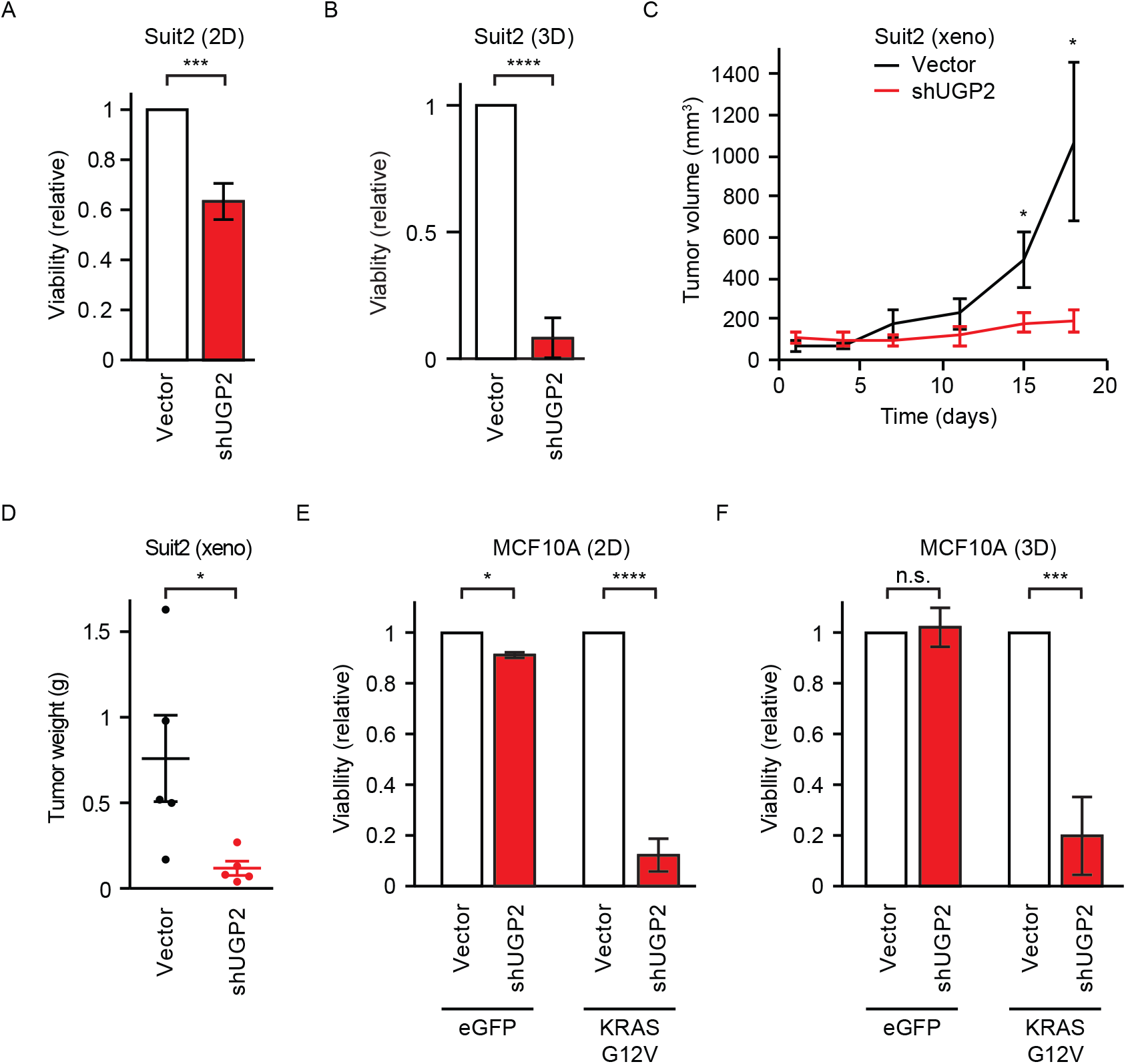
UGP2 is critical for mutant KRAS-driven cell proliferation *in vitro* and *in vivo*. A, Relative viability of Suit2 cells grown in two-dimensional culture conditions for 48 hours, measured by CellTiter-Glo. n = 3, *** p < 0.001. B, Relative viability of Suit2 cells grown in three-dimensional culture conditions for 14 days, n = 4, **** p < 0.0001. C, Tumor volumes of Suit2 cells with shUGP2 or empty vector control xenografted on opposite flanks of nude mice, n = 5, * p < 0.05. D, Tumor weights at endpoint, n = 5, * p < 0.05. E-F, Relative viability of MCF10A cells stably expressing KRAS G12V or control eGFP with shUGP2 or empty vector grown in two-dimensional (E) or three-dimensional matrigel (F) culture conditions. Error bars represent standard error of the mean, n = 3. * p < 0.05, *** p < 0.001, **** p < 0.0001.

To test the extent to which UGP2 was required for tumor growth *in vivo*, PDAC cell lines with stable knockdown of UGP2 or empty vector controls were xenografted onto the opposite flanks of nude mice (Fig. 1C-D and Fig. S1H-J). UGP2 knockdown halted tumor growth in both Suit2 xenografts and MiaPaca2 xenografts over the course of several weeks. Tumors with shUGP2 expressed less UGP2 and displayed a reduced endpoint tumor size. The proliferative index shown by Ki67 staining was decreased in tumors with UGP2 knockdown.

To determine whether UGP2 dependence is limited to oncogenic contexts, UGP2 was knocked down in both control non-oncogenic MCF10A cells and in MCF10A cells transformed by overexpression of oncogenic KRAS G12V, a driver mutation common in PDAC^12^. MCF10A cells transformed with KRAS G12V were highly dependent on UGP2 for proliferation, but knockdown of UGP2 in control MCF10A cells did not result in growth inhibition (Fig. 1E-F, Fig. S1K). Similar results were observed in MCF10A cells grown in both 2D and 3D culture conditions. Together, these data suggest that while UGP2 required for the proliferative growth of cancer cells such as PDAC lines driven by mutant KRAS, it is not universally required for the growth of all cells in culture.

### UGP2 is transcriptionally regulated by the YAP/TEAD complex

Pursuing these indications that UGP2 expression is required for PDAC growth and maintenance, we sought to understand how UGP2 expression is regulated in these contexts. KRAS is mutated in over 90% of PDACs. We found that enforced overexpression of oncogenic KRAS mutants drove an increase in UGP2 mRNA and protein expression (Fig. S2A-C). In addition to the canonical RAF-MEK-ERK effector pathway, KRAS has also been reported to regulate other cellular processes, including programs of largely MAPK-independent metabolic reprogramming^13^. Inhibition of MEK with trametinib did not rescue the increased expression of UGP2 driven by KRAS G12V, leading to the hypothesis that regulation of UGP2 expression is independent of the RAF-MEK-ERK signaling axis (Fig. S2C).

YAP has emerged as a regulator of growth and progression of PDACs by acting as a key transcriptional modulator of its target genes^14,15^. In some murine models, YAP is required for maintenance of pancreatic tumors driven by KRAS G12D^15^. YAP overexpression can recapitulate many of the phenotypes of mutant KRAS-driven cancer and can drive relapse in tumors depleted of oncogenic KRAS^14,16^. Much like UGP2, YAP expression is a negative prognostic marker in PDAC (Fig. S2D). To examine the relationship between YAP and UGP2 in PDACs, immunohistochemistry was employed to probe a tissue microarray panel of 78 PDAC patient samples. Blinded scoring of YAP and UGP2 protein expression in these tumors showed that their expression was strongly positively correlated (p<0.0001) (Fig. 2A and Fig. S2E). Furthermore, the mRNA expression levels of YAP and UGP2 were positively correlated in a panel of 178 PDAC patient samples from the Cancer Genome Atlas (p = 3 × 10^−9^) (Fig. S2F).

**Figure 2.**
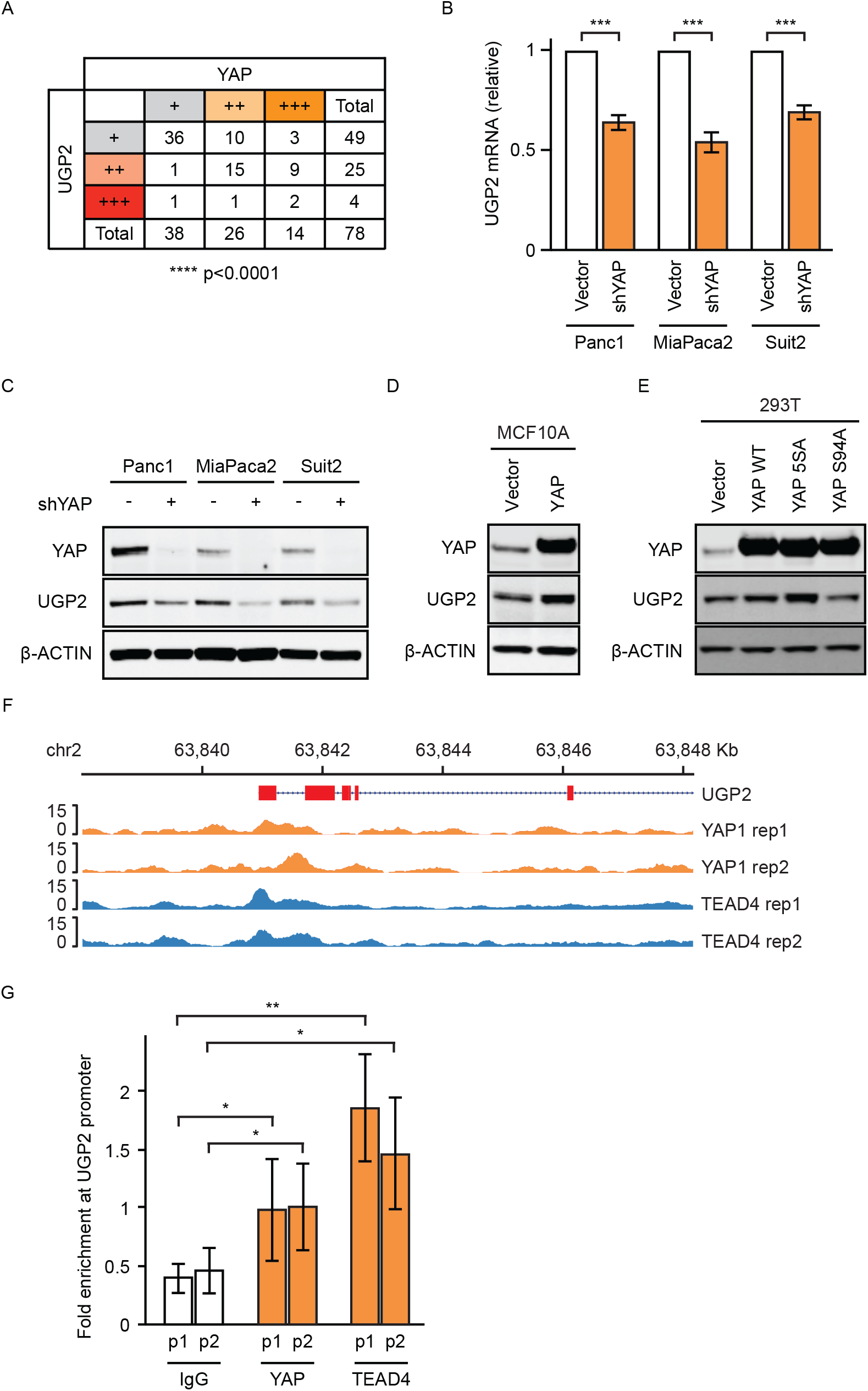
YAP1 directly regulates transcription of UGP2. A, Scoring of tissue microarrays containing 78 pancreatic ductal adenocarcinoma samples immunohistochemically stained for YAP or UGP2. +, ++, +++ represent combined scoring of automated area x intensity measurements. **** p < 0.0001 by Chi-square test. B, qPCR for UGP2 messenger RNA in Panc1, MiaPaca2, and Suit2 cells with shYAP or empty vector. *** p < 0.001. C, Immunoblot on lysates from Panc1, MiaPaca2, and Suit2 cells stably expressing shYAP or empty vector and probed as indicated. D-E, Immunoblots on lysates from MCF10A (D) and 293T (E) cells stably expressing wild-type YAP, YAP5SA, YAP S94A, or empty vector, probed as indicated. F, ChIP-seq for YAP1 and TEAD4 at the UGP2 locus. G, Targeted PCR using two sets of primers (p1 and p2) in the UGP2 promoter region of Panc1 cells after crosslinking and pulldown using antibodies against either YAP, TEAD4, or control IgG. Fold enrichment is percent input target gene/percent input Actin. * p < 0.05, ** p < 0.01.

Given these intriguing correlations, we pursued mechanistic studies to directly test whether YAP regulates the expression of UGP2. Knockdown of YAP in a panel of PDAC cell lines resulted in decreased UGP2 mRNA by qPCR, reduced UGP2 protein by immunoblot, and reduced cellular proliferation (Fig. 2B-C, Fig. S2G-K). Conversely, overexpression of YAP increased UGP2 expression in multiple cell systems (Fig. 2D-E). Introduction of the constitutively active YAP mutant 5SA, in which five inhibitory phosphorylation sites on YAP have been removed^17^, further increased UGP2 expression above that which was observed for wildtype YAP (Fig. 2E). In contrast, expression of the inactivating dominant negative YAP mutant S94A^18^ resulted in a suppression of UGP2 expression.

YAP is a transcriptional co-activator and requires DNA binding partners, such as TEAD4, for transcriptional activation^19,20^. To test whether the YAP/TEAD complex regulates UGP2 transcriptional expression by directly binding at the *UGP2* locus, chromatin immunoprecipitation followed by sequencing (ChIP-seq) was performed for YAP and TEAD4 (Fig. 2F, Fig. S2L). As expected, high occupancy of YAP/TEAD4 was observed at the promoters of well-established YAP/TEAD binding sites including cellular communication network factor 2 (CCN2)^18,19^ (Fig. S2L). Interestingly, we observed binding of both YAP and TEAD4 at the *UGP2* promoter region across biological replicates (Fig. 2F). Specific binding of YAP and TEAD4 was confirmed using ChIP-qPCR primer pairs targeted to the UGP2 promoter (Fig. 2G). These converging lines of evidence support a model wherein transcription of UGP2 is positively regulated by direct binding of the YAP/TEAD complex to the UGP2 locus.

### UGP2 regulates protein glycosylation in PDAC cells

UGP2 enzymatic activity is the rate-limiting step in UDP-glucose production. This activated form of glucose is a substrate of the N-linked glycosylation pathway^21^, however the specific modifications that UGP2 regulates in cancer cells have not been elucidated. We performed unbiased global N-glycoproteomic analysis of all asparagine-linked glycan modifications in Suit2 cells following 48 hours of UGP2 knockdown using siRNA (Fig. 3A, Fig. S3A). A large-scale decrease in glycan modifications was observed relative to cells treated with scrambled siRNA controls. Knockdown of UGP2 significantly decreased the incidence of 141 N-glycosylation modifications spread across 89 proteins (Table S1). Total global proteomics, performed in parallel, revealed that the vast majority of the observed decreases in N-glycosylation were not explained by corresponding changes in total protein levels or global changes in the relative frequency of glycan chain architectures (Fig. S3B-D, Table S1). Thus, UGP2 knockdown in PDAC cells rapidly reduces N-glycosylation modifications in a distinctive constellation of sites across multiple proteins.

**Figure 3.**
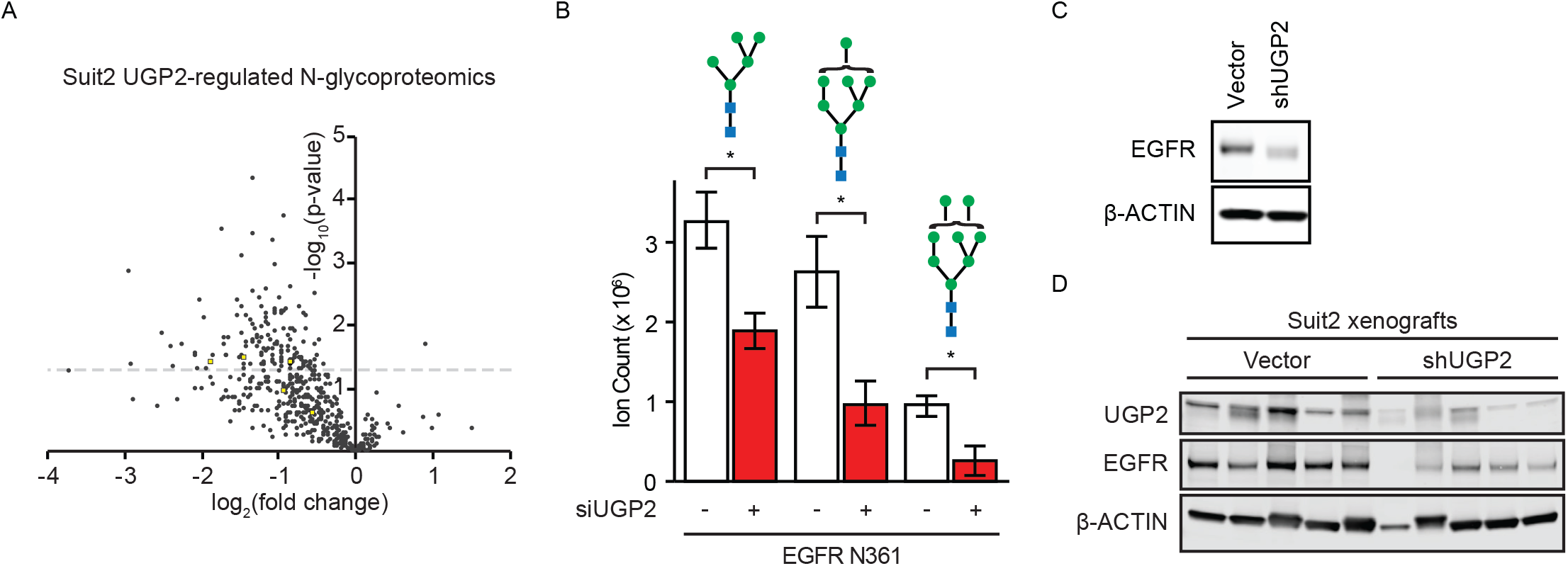
UGP2 regulates N-glycan modifications on proteins including EGFR. A, Unbiased quantitative N-glycoproteomic analysis identified changes in protein modifications in Suit2 cells upon knockdown of UGP2 relative to non-targeting control siRNAs at 48 hours, yellow boxes represent modifications on EGFR. B, Quantification of glycan modifications on EGFR Asn361 in Suit2 cells with siUGP2 or non-targeting control siRNAs at 48 hours. * p < 0.05, blue squares represent N-acetylglucosamines, green circles represent mannoses. C, Immunoblot of lysates from Panc1 cells with shUGP2 or empty vector, probed as indicated. D, Immunoblot of Suit2 xenograft endpoint tumor lysates, probed as indicated.

Of the 89 proteins with N-glycan modifications significantly regulated by UGP2, EGFR was among the top proteins whose N-glycan modifications were most frequently and consistently decreased upon knockdown of UGP2 (Fig. 3B). EGFR is a membrane-embedded growth factor receptor whose activation leads to signaling through the RAS-MEK-ERK growth axis, upon which many PDACs are dependent. Mutations and genomic amplifications in EGFR are associated with a wide variety of oncogenic malignancies and N-glycosylation of EGFR is required for its trafficking, efficient ligand binding, and receptor activation^22^. We identified an UGP2-regulated N-glycosylation site on the extracellular ligand binding domain of EGFR at Asn361, where three different N-glycan modifications were significantly decreased by knockdown of UGP2 (Fig. 3A-B). Decrease of an additional N-glycan modification to EGFR was observed at Asn528, although this change did not reach statistical significance (Fig. S3E). While the amount of total EGFR protein remained stable upon transient 48-hour siRNA-based perturbation of UGP2 (Fig. S3F), immunoblots for EGFR in cells with long-term stable shRNA knockdown of UGP2 showed a decrease in total EGFR protein and a gel shift in EGFR size (Fig. 3C-D, Fig. S3G). In the majority of xenografted tumors, knockdown of UGP2 resulted in decreased EGFR protein as shown in both immunoblots of tumor lysates and immunohistochemistry on tumor sections (Fig. S1J). Hence, UGP2 loss leads to decreases in EGFR N-glycosylation at Asn361 and ultimately a decrease in total EGFR protein.

### Effects of UGP2 loss on glycogen synthesis

Another important cellular process that utilizes UDP-glucose is glycogen synthesis^23^. Glycogen is a multibranched polysaccharide that serves as an important form of energy for proliferating cancer cells, especially in glucose-depleted conditions^24^. To investigate the effect of UGP2 on cell survival in response to glucose starvation, we assessed the viability of UGP2-depleted Panc1 cells in standard tissue culture media with 25 mM glucose or reduced glucose concentrations of 2.5 mM, 0.25 mM, or 0 mM. Knockdown of UGP2 caused a stronger growth-inhibitory effect in the low glucose conditions (Fig. 4A). Furthermore, UGP2 knockdown-induced growth inhibition in glucose-starved cells was rescued by supplement with its enzymatic product UDP-glucose, whereas adding UDP-glucose had no effect on control cells (Fig. 4B, Fig. S4A). These data suggest that loss of UGP2 creates a scarcity of UDP-glucose that inhibits survival in low-nutrient contexts.

**Figure 4.**
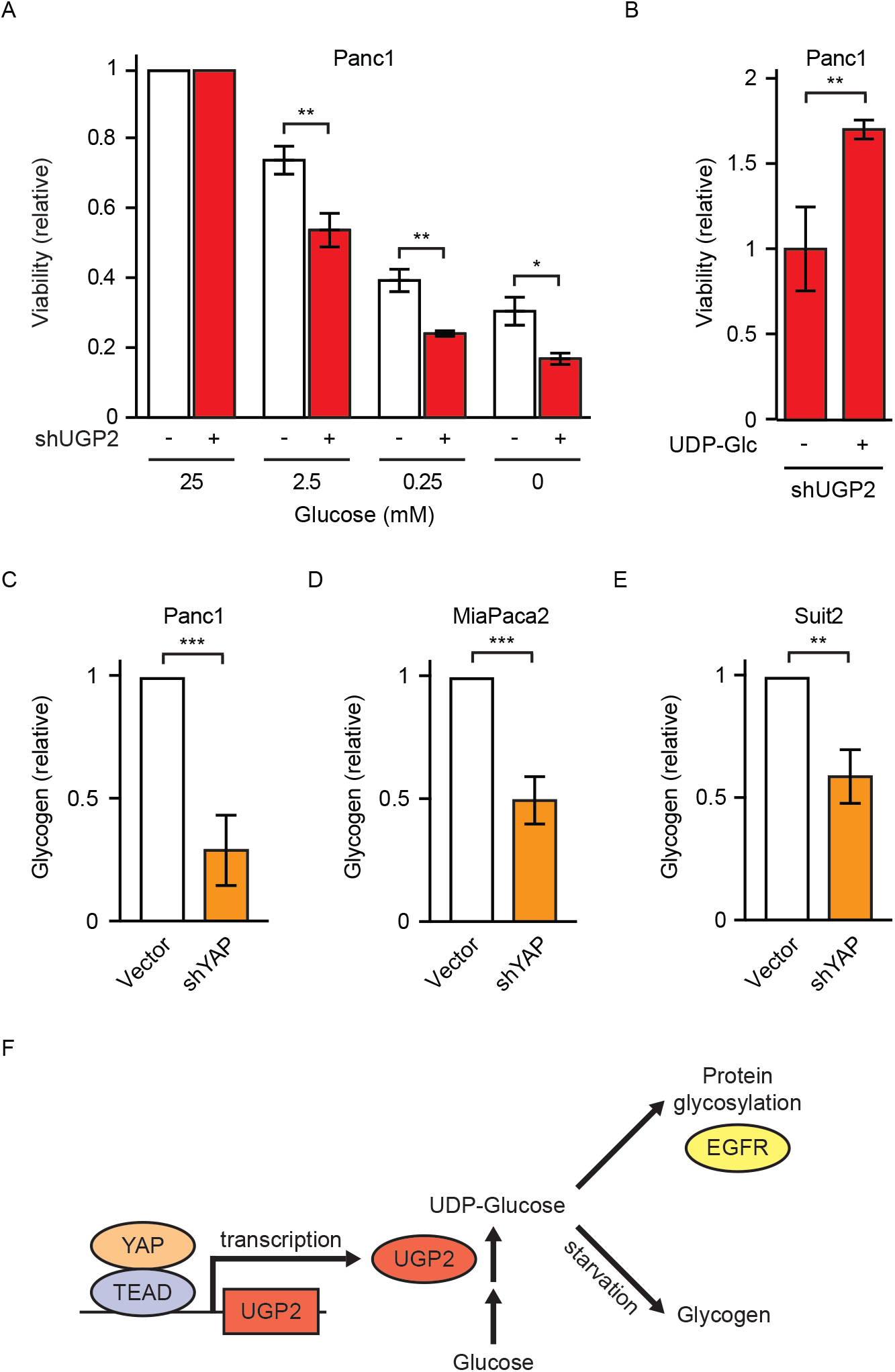
Effects of UGP2 loss on cellular processes. A, Viability relative to complete media of Panc1 cells with shUGP2 or empty vector in the indicated concentration of glucose, * p < 0.05, ** p < 0.01. B, Relative viability of Panc1 cells with shUGP2 with or without 100 μM UDP-Glc for 7 days, ** p < 0.01. C-E, Cellular glycogen in Panc1 (C), MiaPaca2 (D), or Suit2 (E) cells with shYAP or empty vector control. ** p < 0.01, *** p < 0.001, error bars represent SEM. F, Model of UGP2 action in cancer.

Finally, we investigated upstream regulation of UGP2 and its effects on glycogen synthesis. Knockdown of YAP decreased cellular UDP-glucose, consistent with a model wherein YAP tonically positively regulates the expression of UGP2 (Fig. S4B-C). Knockdown of YAP led to depletion of cellular glycogen in three PDAC cell lines (Fig. 4C-E), and conversely YAP overexpression led to increased cellular glycogen (Fig. S4D). Thus, YAP regulates the production of UDP-glucose and regulates glycogen production, processes which are also regulated by UGP2. UGP2 plays an important role in the metabolism of PDAC cells by regulating protein glycosylation as well as glycogen synthesis (Fig. 4F).

## Discussion

PDACs are extremely intractable solid tumors that experience a hypoxic and nutrient-deprived microenvironment and often undergo a metabolic switch to support increased glycolytic flux for their growth^25^. These differential metabolic settings may provide novel opportunities to selectively target cancers via their metabolism. For example, L-asparaginase, an enzyme that induces asparagine deprivation, has been successfully used to treat acute lymphoblastic leukemia and is now undergoing clinical trials for PDAC^25^. Employing *in vitro* 2D and 3D models as well as *in vivo* tumor xenograft models, we identified UGP2 as a critical regulator of protein N-glycosylation and glycogen synthesis that is essential for PDAC cell growth, exposing a vulnerability that could be exploited for therapeutic interventions. An intriguing question is how ubiquitous UGP2 dependency is across different cancer types. In the Dependency Map (DepMap) portal, cell lines from multiple tissue types display a dependency on UGP2 for their growth^26,27^, suggesting that the requirement for UGP2 may extend beyond PDACs and other mutant KRAS driven malignancies. Interestingly, UGP2 is co-dependent with multiple other genes which have a role in protein glycosylation including phosphoglucomutase 3 (PGM3), phosphomannomutase 2 (PMM2), and mannose phosphate isomerase (MPI), highlighting the biological complexity of this pathway.

We identified YAP as an important new transcriptional regulator of UGP2 expression in PDAC. YAP is required for maintenance of some KRAS mutant tumors and in a relapse setting activated YAP mutants can largely phenocopy KRAS to drive PDAC even in the absence of KRAS^14^. A key function of YAP is to act as a transcriptional modulator regulating metabolic processes, however the critical effectors of YAP in regulating metabolism remained elusive. Here we identify UGP2 as a key effector of YAP function to regulate growth in PDAC cells, providing a potential mechanism for how increased YAP activity observed in cancer may drive increases in tumor metabolism and subsequent growth.

Unbiased global N-glycoproteomics identified 89 proteins with N-glycosylation sites significantly reduced by knockdown of UGP2 in PDAC cells and understanding which of these post-translational modifications are functionally significant remains an important step for subsequent studies. UGP2 loss led to decreased glycosylation of several proteins with well-established roles in PDAC biology, such as c-MET and EGFR (Table S1). Depletion of UGP2 also led to a subsequent decrease in total EGFR protein, which Suit2 cells depend upon for growth^26,27^, although the mechanism by which this decrease occurred remains unclear. Differential glycosylation can drive changes in protein behaviors including degradation, localization, and signaling and can act as a diagnostic marker^7^. Glycan modifications may be important for promoting EGFR stability or inhibiting its degradation. The proximity of Asn361 to the ligand-binding site suggests that the UGP2-regulated family of EGFR modifications could affect the ability of EGF ligands to bind and initiate downstream signaling cascades. The role of UGP2 in cancer progression *in vivo* warrants further investigation as an intrinsic mechanism and as a node for discovery of novel pharmacological interventions.

## Materials and Methods

### Tissue culture and xenografts

All cell lines were confirmed by STR analysis and were regularly screened for mycoplasma. Panc1, MiaPaca2, Suit2, and HEK293T were grown in DMEM (Invitrogen) with 10% FBS (Atlanta Biologicals). MCF10A cells were grown in DMEM/F12 (Invitrogen) supplemented with 5% horse serum (Invitrogen), 20 ng/mL EGF (Peprotech), 0.5 mg/mL hydrocortisone (Sigma), 100 ng/mL cholera toxin (Sigma), and 10 μg/mL insulin (Sigma). Prior to experiments, 2 μg/mL puromycin (Invitrogen A11138-02) was used to select for eGFP, KRAS G12D, or G12V. 4 μg/mL blasticidin (Fisher AAJ61883WDL) was used to select for the shRNA of interest or the empty vector pLKO. 3D cultures were seeded using 20% matrigel (Corning 354234) on low attachment 96-well plates (Corning 3474) for 2 weeks.

Cell viability was measured using CellTiter-Glo luminescent cell viability assay (Promega G7570). Cellular glycogen was measured using the glycogen assay kit (Abcam ab65620). Metabolomics including UDP-glucose levels were measured by Precision Metabolomics LC-MS global metabolomics platform (Metabolon).

For each cell line, five nude mice (NCr-Foxn1nu from Taconic) were injected with 3,000,000 cells expressing either LKO or shUGP2 resuspended in 70% PBS/30% matrigel on opposite flanks. Subsequent tumors were monitored by caliper 2-3 times per week beginning one day after injection. Sample sizes required for statistically meaningful results were determined based on standards in the field and minimization of animal use. All animal studies and endpoints were consistent with UCSF institutional animal care and use committee guidelines. At endpoint, tumors were weighed and cut into portions that were snap-frozen for immunoblots or fixed for histology using 24 hours of immersion in 10% neutral buffered formalin. Tumor lysates were prepared using a Fisher Powergen Homogenizer for 30-60 seconds in cell lysis buffer (20 mM Tris pH 7.4, 150 mM NaCl, 1% NP-40, 1 mM EDTA, 1 mM EGTA, 10% glycerol, 50 mM NaF, 5 mM NaPPl, 1 mM PMSF, 0.5 mM DTT, PPase inhibitor, protease inhibitor). Lysates were incubated on ice for 30 minutes and spun at 13,000 rpm at 4°C, then the supernatant was frozen at −80°C.

### Immunoblots and immunohistochemistry

Lysates were diluted with 4X NuPage buffer and run on Invitrogen NuPage 4-12% Bis-Tris gels. The antibodies employed were KRAS (Sigma 3B10-2F2), UGP2 (Santa Cruz Biotechnology sc-377089), YAP (Cell Signaling 14074), EGFR (Cell Signaling 4267), and B-ACTIN (Sigma A5441).

Immunohistochemistry of xenografts was performed by Histowiz using antibodies against EGFR (EP38Y), and UGP2 (Santa Cruz Biotechnology sc-377089 at 1:200). Immunohistochemistry of historic deidentified PDAC samples used antibodies against YAP1 (Cell Signaling 14074) and UGP2 (Santa Cruz Biotechnology sc-377089). To analyze tissue microarray staining, blinded data extraction for mean-intensity and percent-area was conducted using inForm software from PerkinElmer. Normalization and Chi-square statistical analysis were conducted using Matlab.

### RNA interference

shRNAs targeting YAP and UGP2 were purchased from Sigma. The shRNA constructs were packaged as lentiviruses using third generation packaging systems with standard protocols. The YAP shRNA sequences were: CCCAGTTAAATGTTCACCAAT (TRCN0000107265) and GCCACCAAGCTAGATAAAGAA (TRCN0000107266). The UGP2 shRNA sequences were: CCACAGCATCATCACATGAAT (TRCN0000037840) and GCTAGTTTCTTACAATGAAAT (TRCN0000435330).

For siRNA experiments, Suit2 cells were grown in 2D culture in 15 cm plates (Corning 430599) and were transfected with pools of non-targeting siCtrl or siUGP2. siRNAs in the control non-targeting pool (D-001810-10-05) and UGP2 ON-TARGETplus SMARTpool (L-007739-00-0050) were purchased from Dharmacon. The UGP2 siRNA sequences in the pool were: GAGCUAGAAUUAUCUGUGA, UAGCAAAGGACGUGUCUUA, ACAAACAACCUAUGGAUUU, and UAAUAUAUCUUCCGUGUUG. Transfections were performed using DharmaFECT 1 Transfection Reagents (Dharmacon T-2001). After 48 hours, triplicate cell pellets were washed twice in PBS, harvested, counted, pelleted, and frozen in ~10,000,000 aliquots for downstream immunoblot, N-glycomic, N-glycoproteomic, and total proteomic analyses.

### Chromatin immunoprecipitation

5,000,000 cells were seeded in 10 cm plates in triplicates. Crosslinking was performed by adding 37% (wt/vol) formaldehyde to a final concentration of 1.42% drop-wise to the plate at room temperature. Cells were incubated with rotation for 15 minutes at room temperature for 15 minutes. Then the formaldehyde was quenched with ice cold 125 mM glycine for 5 minutes at room temperature. Cells were scraped, washed twice in ice cold PBS, and frozen on dry ice. Pellets were resuspended in 1 mL ChIP buffer (330 μL Triton buffer + 660 μL SDS buffer + 10 μL protease inhibitor). Global ChIP-seq and UGP2 promoter ChIP-PCR were performed using antibodies against YAP1 (Novus Biologicals NB110-58358) and TEAD4 (ab58310). ChIP-PCR was performed with two independent sets of PCR primers: UGP2 (2-1) Forward primer AGAGGTTGGTGGGTGGTTTG Reverse primer ATACGCGTCTGGAACGTCA UGP2 (3-1) Forward primer TTGTGTGTACGTGGTTTGCG Reverse primer AATGACCTCCAGCTTCTTCGG.

### Glycomics, glycoproteomics, and total proteomics

Cell membrane extraction methods were applied as previously described with slight modifications^28–30^. Cell pellets were resuspended in a homogenization buffer containing 20 mM HEPES (pH 7.4), 0.25 M sucrose, and a 1:100 protease inhibitors (Calbiochem/EMD Chemicals). Cells were lysed with five alternating on and off pulses in 5 and 10 second intervals using a probe sonicator (Qsonica) followed by centrifugation at 2000 × g for 10 minutes and ultracentrifugation at 200,000 x g for 45 minutes at 4°C. Pellets were resuspended in 500 μL of 0.2 M Na_2_CO_3_ and 500 μL of water followed by two additional ultracentrifugations at 200,000 × g for 45 minutes at 4°C.

For protein digestion, the cell membrane fractions were reconstituted in 60 μL of 8M urea and sonicated for 10 minutes to homogenize the pellet^28–30^. To further the denaturing process, 2 μL of dithiothreitol (DTT) was added to each sample and incubated for 50 minutes at 55°C, followed by the addition of 4 μL of iodoacetamide (IAA) for 20 minutes at room temperature in the dark. 420 μL ammonium bicarbonate (NH_4_HCO_3_) solution was added to adjust the pH and 2 μg of trypsin for protein digestion was added for 18 hours at 37°C. Peptides were desalted by solid-phase extraction (SPE) with C18 cartridges (Sigma) containing 500 mg materials for proteomic analysis or enriched by SPE using iSPE®-HILIC cartridges (The Nest Group) for glycopeptide analysis. Samples were dried in vacuo using miVac (SP Scientific).

Glycopeptide samples were reconstituted in 10 μL of nanopure water prior to injection into a Nanospray Flex ion source Orbitrap Fusion Lumos Tribrid Mass spectrometer coupled with an UltiMate™ WPS-3000RS nanoLC 980 system (Thermo Fisher Scientific) conducted with a binary gradient system where solvent A composed of 0.1% (v/v) formic acid and solvent B composed of 80% (v/v) acetonitrile^31^. Samples were separated on an Acclaim™ PepMap™ 100 C18 LC Column (3 μm, 0.075 mm × 150 mm, Thermo Fisher Scientific) with the following gradient sequence for separation: 0–5 min, 4–4% (B); 5–133 min, 4–32% (B); 133–152 min, 32%-48% (B); 152–155 min, 48–100% (B); 155–170 min, 100–100% (B); 170–171 min, 100–4% (B); 171–180 min, 4–4% (B). Data acquisition was performed with mass range of m/z 700 to 2000 in positive ionization mode. Ionization spray voltage was set to 1.8 kV with a 275°C ion transfer capillary temperature. Precursor ions were subjected to stepped higher-energy C-trap dissociation (30±10%) applied to obtain tandem MS/MS spectra with a minimum mass range set at 120 m/z. Proteomic analysis was reconstituted to 60 μL and was performed on the same instrument with the following gradient sequence for separation: 0–5 min, 4– 4% (B); 5–90 min, 4–47% (B); 90–100 min, 47%-70% (B); 100–100.5 min, 70–100% (B); 100.5–115.5 min, 100–100% (B); 115.5–116 min, 100–4% (B); 116–130 min, 4–4% (B).

Raw files from the glycoproteomic and proteomic analysis were identified from a Uniprot FASTA database using Byonic and Byologic software (Protein Metrics)^31^. Search parameters were set with precursor mass tolerance of 10 ppm and QTOF/HCD fragmentation with mass tolerance of 20 ppm. Fixed modification was assigned to carbamidomethylation at cysteine and variable modification of deamidated amino acid were assigned to asparagine and glutamine, methylation of lysine and arginine.

Cell membrane fractions were reconstituted in 100 μL of 100 mM NH_4_HCO_3_ with 5 mM of DTT^32^. Samples were heated for 1 minute at 100°C followed by the addition of 2 μL of peptide N-glycosidase F (New England Biolabs) to release the N-glycans. Samples were then incubated in a microwave reactor (CEM Corporation) at 20 watts, 37°C for 10 min. The resulting solutions were incubated in a 37°C water bath for 18 hours. Ultracentrifugation at 200,000 × g for 45 minutes was performed to isolate the N-glycans fractions from the protein precipitates. N-glycans were purified using a porous graphitic carbon (PGC) 96 well SPE plate. Samples were dried in vacuo using miVac (SP Scientific).

N-glycan samples were reconstituted in 60 μL of nanopure water prior to injection in Agilent 6520 Accurate Mass Q-TOF LC/MS coupled with a PGC nano-chip (Agilent Technologies). N-glycan separation was conducted with a binary solvent system where solvent A was composed of 0.1% (v/v) formic acid and 3% (v/v) acetonitrile and solvent B was composed of 1% (v/v) formic acid and 90% (v/v) acetonitrile. The gradient sequence for separation was: 0–2 min, 3–3% (B); 2–20 min, 3–16% (B); 20–40 min, 16%-72% (B); 40–42 min, 72–100% (B); 42–52 min, 100–100% (B); 52–54 min, 100–0% (B); 54–65 min, 3–3% (B) with a constant flow rate of 300 nL min-1. Spectra were collected every 1.5 seconds in positive mode ionization with a mass range of m/z 600-2000. The top five most abundant precursor ions in each MS1 spectrum were subjected to collision-induced dissociation (CID) fragmentation. The collision equation is Vcollision = 1.8 × (m/z) /100 V −2.4 V. Data were analyzed using MassHunter Qualitative Analysis B08 software (Agilent) Find by Molecular Feature function with mass tolerance of 20 ppm. N-glycans were identified based on an in-house library with accurate mass. Relative abundances were determined based on integrated peak areas for each glycan composition and normalized to the summed peak areas of all detected glycans.

## Acknowledgements

We thank the UCSF Biorepository & Tissue Biomarker Technology Core for TMA image analysis. Histowiz, Inc. performed immunohistochemical staining of murine tumors and UDP-glucose levels were measured by Metabolon, Inc. Patient PDAC mRNA expression data was generated by The Cancer Genome Atlas Research Network. We thank members of the McCormick lab for discussions and critical reading of the manuscript. Funding for this work was provided by Damon Runyon Cancer Research Foundation DRG-2214-15 (A.L.W.), NIH R01GM049077 (Q.Z., C.B.L.), MSKCC Support Grant/Core Grant P30 CA008748 (M.S.), and the UCSF Research Allocation Program (S.E.K.). Research reported in this publication was supported by the National Cancer Institute of the National Institutes of Health under Award Number K99CA226363 (A.L.W.). The content is solely the responsibility of the authors and does not necessarily represent the official views of the National Institutes of Health.

## Author Contributions

A.L.W. and S.E.K. designed and executed experiments and prepared the manuscript. Q.Z. and C.B.L. provided expertise, performed experiments, and wrote methods for all proteomic analyses. J.G. provided technical assistance with experiments. E.T., R.T.K., and M.S. performed and analyzed ChIP-Seq experiments. A.A.K. and S.B. provided expression data of UGP2 in MCF10A cells. F.M. provided valuable advice and guidance throughout.

## Competing Interests

M.S. has received research funds from Puma Biotechnology, AstraZeneca, Daiichi Sankyo, Immunomedics, Targimmune and Menarini Ricerche. He is in the scientific advisory board of Menarini Ricerche and Bioscience Institute and is a cofounder of medendi.org. F.M. is a consultant for Daiichi-Sankyo, Pfizer, Amgen and has received research funding from Daiichi Sankyo. The other authors declare no potential conflicts of interest.

## Supplemental figure legends

**Figure S1.** UGP2 is critical for cell growth in cells with mutant Ras.

A, Cumulative probability of survival of 177 patients with pancreatic ductal adenocarcinoma split by expression of UGP2 (p = 0.016). Data adapted from cBioPortal ^33,34^. B, Immunoblots for shRNAs knocking down UGP2 in Panc1 cells, probed as indicated. C-D, Relative viability by CellTiter-Glo after 48 hours of growth in 2D culture of MiaPaca2 (C) and Suit2 (D) cells with shUGP2 or empty vector control. **p<0.01, ***p<0.001. E-F, Representative crystal violet staining of Suit2 (E) and Panc1 (F) cells with two independent shRNAs against UGP2 or empty vector control grown in two-dimensional culture for 10 days. G, Representative images of Panc1 and Suit2 cells stably expressing shUGP2 or Vector control grown in three-dimensional matrigel for 14 days. H, Tumor volumes of MiaPaca2 cells with shUGP2 or empty vector control xenografted on opposite flanks of nude mice, n = 4, * p < 0.05. I, Tumor weights at endpoint, n = 4, * p < 0.05. J, Representative immunohistochemical staining of paired opposite-flank tumors with shUGP2 or vector control at endpoint, stained as indicated. Scale bar represents 50 μM. K, Representative images of MCF10A cells stably expressing shUGP2 or empty vector grown in 2D or 3D matrigel for 14 days, n = 3.

**Figure S2.** Regulation of UGP2 by YAP1.

A, mRNA expression of UGP2 from RNAseq on MCF10A cells stably expressing KRAS G12V or empty vector. B-C, Immunoblots on lysates from MCF10A cells stably expressing KRAS G12D, KRAS G12V, or empty vector (B) or treated with 100 nM trametinib for 48 hours (C). D, Cumulative probability of survival in PDAC patient samples split by expression of YAP1, n = 177, *** p = 0.00021. E, Representative tissue microarray images stained for YAP or UGP2. B-9, B-15, and G-11 indicate sample codes. F, Correlation between UGP2 and YAP1 mRNA in pancreatic ductal adenocarcinoma patient samples. Analysis performed using cBioPortal, n = 178, R^2^ = 0.18, Pearson = 0.43, p = 3*10-9 ^33,34^. G-I, Immunoblots on cells with shRNAs against YAP or vector control, probed as indicated. Shown are Panc1 with four different YAP shRNAs (G), MCF10A stably expressing KRAS G12V or control eGFP (H), and a panel of pancreatic cancer cell lines (I). J-K, Relative viability of MiaPaca2 (J) and Suit2 (K) cells by CellTiter-Glo after 48 hours of growth in two-dimensional culture in with shYAP or empty vector control. L, ChIP-seq for the control YAP1/TEAD4 binding site in the CCN2 genomic region.

**Figure S3.** Glycan changes induced by UGP2 knockdown.

A, Immunoblots on lysates from 293T or Suit2 cells transfected with siUGP2 or non-targeting control siRNAs for 48 hours, probed as indicated. B, Global total proteomic analysis in Suit2 cells upon knockdown of UGP2 relative to non-targeting control siRNAs after 48 hours, yellow boxes represent total EGFR. C, Shown are log2 of the fold change in specific modifications plotted against log2 of the fold change in total protein upon knockdown of UGP2 in Suit2. R^2^ = 0.0215, yellow boxes represent EGFR modifications. D, Global glycomic comparison in Suit2 cells with siUGP2 of non-targeting control siRNAs at 48 hours, n.s. not significant at a threshold of p < 0.05. E, Quantification of glycan modifications on EGFR N528 in Suit2 cells with siUGP2 or non-targeting control siRNAs at 48 hours. n.s. not significant at a threshold of p < 0.05, blue squares represent N-acetylglucosamines, green circles represent mannoses, yellow circles represent galactoses, red triangles represent fucoses, and purple diamonds represent N-acetylneuraminic acid. F, Quantification of total EGFR, α-TUBULIN, and GAPDH proteins by mass spectrometry in Suit2 cells upon knockdown of UGP2 or non-targeting control siRNAs. n.s. not significant at a threshold of p < 0.05. G, Immunoblot of MiaPaca2 xenograft endpoint tumor lysates, probed as indicated.

**Figure S4.** Functional effects of UGP2 and YAP perturbation.

A, Relative viability of Panc1 cells with vector control grown with or without 100 μM UDP-Glc for 7 days, n.s. not significant at a threshold of p < 0.05. B, Global metabolomics of Panc1 cells with shYAP or empty vector control. Orange dots indicate metabolites in shYAP cells significantly decreased p < 0.05, blue dots indicate metabolites significantly increased p < 0.05, arrow indicates UDP-Glucose. C, Relative glycogen in MCF10A cells expressing YAP or empty vector control. ** p <0.01. D, UDP-glucose levels or Panc1, MiaPaca2, and Suit2 cells with shYAP or empty vector. * p < 0.05 by ANOVA, n.s. not significant.

**Table S1.** N-glycoproteomics and total proteomics.

A-B, Changes in N-glycan modifications (A) and total protein (B) in Suit2 cells treated with siUGP2 or siCtrl pools for 48 hours, n = 3.

## References

1. Ying, H. et al. Genetics and biology of pancreatic ductal adenocarcinoma. Genes Dev. 30, 355–385 (2016).

2. Commisso, C. et al. Macropinocytosis of protein is an amino acid supply route in Ras-transformed cells. Nature 497, 633–637 (2013).

3. Kamphorst, J. J. et al. Human Pancreatic Cancer Tumors Are Nutrient Poor and Tumor Cells Actively Scavenge Extracellular Protein. Cancer Research 75, 544–553 (2015).

4. Grasso, C., Jansen, G. & Giovannetti, E. Drug resistance in pancreatic cancer: Impact of altered energy metabolism. Critical Reviews in Oncology/Hematology 114, 139–152 (2017).

5. Ying, H. et al. Oncogenic Kras Maintains Pancreatic Tumors through Regulation of Anabolic Glucose Metabolism. Cell 149, 656–670 (2012).

6. Pan, S., Brentnall, T. A. & Chen, R. Glycoproteins and glycoproteomics in pancreatic cancer. World J. Gastroenterol. 22, 9288–9299 (2016).

7. Li, Q. et al. Site-Specific Glycosylation Quantitation of 50 Serum Glycoproteins Enhanced by Predictive Glycopeptidomics for Improved Disease Biomarker Discovery. Anal. Chem. 91, 5433–5445 (2019).

8. Wagatsuma, T. et al. Discovery of Pancreatic Ductal Adenocarcinoma-Related Aberrant Glycosylations: A Multilateral Approach of Lectin Microarray-Based Tissue Glycomic Profiling With Public Transcriptomic Datasets. Front Oncol 10, 338 (2020).

9. Führing, J. I. et al. A Quaternary Mechanism Enables the Complex Biological Functions of Octameric Human UDP-glucose Pyrophosphorylase, a Key Enzyme in Cell Metabolism. Sci Rep 5, 9618 (2015).

10. Wang, L. et al. Expression of UGP2 and CFL1 expression levels in benign and malignant pancreatic lesions and their clinicopathological significance. World J Surg Onc 16, 11 (2018).

11. Wang, X. et al. UDP-glucose accelerates SNAI1 mRNA decay and impairs lung cancer metastasis. Nature (2019) doi:10.1038/s41586-019-1340-y.

12. Konishi, H. et al. Knock-in of Mutant K-ras in Nontumorigenic Human Epithelial Cells as a New Model for Studying K-ras Mediated Transformation. Cancer Research 67, 8460–8467 (2007).

13. Pylayeva-Gupta, Y., Grabocka, E. & Bar-Sagi, D. RAS oncogenes: weaving a tumorigenic web. Nature Reviews Cancer 11, 761–774 (2011).

14. Kapoor, A. et al. Yap1 Activation Enables Bypass of Oncogenic Kras Addiction in Pancreatic Cancer. Cell 158, 185–197 (2014).

15. Zhang, W. et al. Downstream of mutant KRAS, the transcription regulator YAP is essential for neoplastic progression to pancreatic ductal adenocarcinoma. Science signaling 7, ra42 (2014).

16. Shao, D. D. et al. KRAS and YAP1 Converge to Regulate EMT and Tumor Survival. Cell 158, 171–184 (2014).

17. Zhao, B. et al. Inactivation of YAP oncoprotein by the Hippo pathway is involved in cell contact inhibition and tissue growth control. Genes & Development 21, 2747–2761 (2007).

18. Zhao, B. et al. TEAD mediates YAP-dependent gene induction and growth control. Genes & Development 22, 1962–1971 (2008).

19. Stein, C. et al. YAP1 Exerts Its Transcriptional Control via TEAD-Mediated Activation of Enhancers. PLoS Genet 11, e1005465 (2015).

20. Vassilev, A. TEAD/TEF transcription factors utilize the activation domain of YAP65, a Src/Yes-associated protein localized in the cytoplasm. Genes & Development 15, 1229–1241 (2001).

21. Deng, Z., Yi, X., Chu, J. & Zhuang, Y. A study on enhanced O-glycosylation strategy for improved production of recombinant human chorionic gonadotropin in Chinese hamster ovary cells. Journal of Biotechnology 306, 159–168 (2019).

22. Kaszuba, K. et al. N-Glycosylation as determinant of epidermal growth factor receptor conformation in membranes. Proc Natl Acad Sci USA 112, 4334–4339 (2015).

23. Higuita, J.-C., Alape-Girón, A., Thelestam, M. & Katz, A. A point mutation in the UDP-glucose pyrophosphorylase gene results in decreases of UDP-glucose and inactivation of glycogen synthase. Biochemical Journal 370, 995–1001 (2003).

24. Favaro, E. et al. Glucose Utilization via Glycogen Phosphorylase Sustains Proliferation and Prevents Premature Senescence in Cancer Cells. Cell Metabolism 16, 751–764 (2012).

25. Cohen, R. et al. Targeting cancer cell metabolism in pancreatic adenocarcinoma. Oncotarget 6, 16832 (2015).

26. Broad DepMap. DepMap Achilles 18Q3 public. 2485731861 Bytes (2018) doi:10.6084/M9.FIGSHARE.6931364.V1.

27. Meyers, R. M. et al. Computational correction of copy number effect improves specificity of CRISPR–Cas9 essentiality screens in cancer cells. Nat Genet 49, 1779–1784 (2017).

28. An, H. J. et al. Extensive Determination of Glycan Heterogeneity Reveals an Unusual Abundance of High Mannose Glycans in Enriched Plasma Membranes of Human Embryonic Stem Cells. Mol Cell Proteomics 11, M111.010660 (2012).

29. Park, D. et al. Characteristic Changes in Cell Surface Glycosylation Accompany Intestinal Epithelial Cell (IEC) Differentiation: High Mannose Structures Dominate the Cell Surface Glycome of Undifferentiated Enterocytes. Mol Cell Proteomics 14, 2910–2921 (2015).

30. Park, D. D. et al. Membrane glycomics reveal heterogeneity and quantitative distribution of cell surface sialylation. Chem. Sci. 9, 6271–6285 (2018).

31. Li, Q., Xie, Y., Xu, G. & Lebrilla, C. B. Identification of potential sialic acid binding proteins on cell membranes by proximity chemical labeling. Chem. Sci. 10, 6199–6209 (2019).

32. Xu, G. et al. Unveiling the metabolic fate of monosaccharides in cell membranes with glycomic and glycoproteomic analyses. Chem. Sci. 10, 6992–7002 (2019).

33. Cerami, E. et al. The cBio Cancer Genomics Portal: An Open Platform for Exploring Multidimensional Cancer Genomics Data: Figure 1. Cancer Discovery 2, 401–404 (2012).

34. Gao, J. et al. Integrative Analysis of Complex Cancer Genomics and Clinical Profiles Using the cBioPortal. Science Signaling 6, pl1–pl1 (2013).

