## Supplemental Figures S1-S4 for "UDP-glucose pyrophosphorylase 2, a regulator of glycosylation and glycogen, is essential for pancreatic cancer growth"

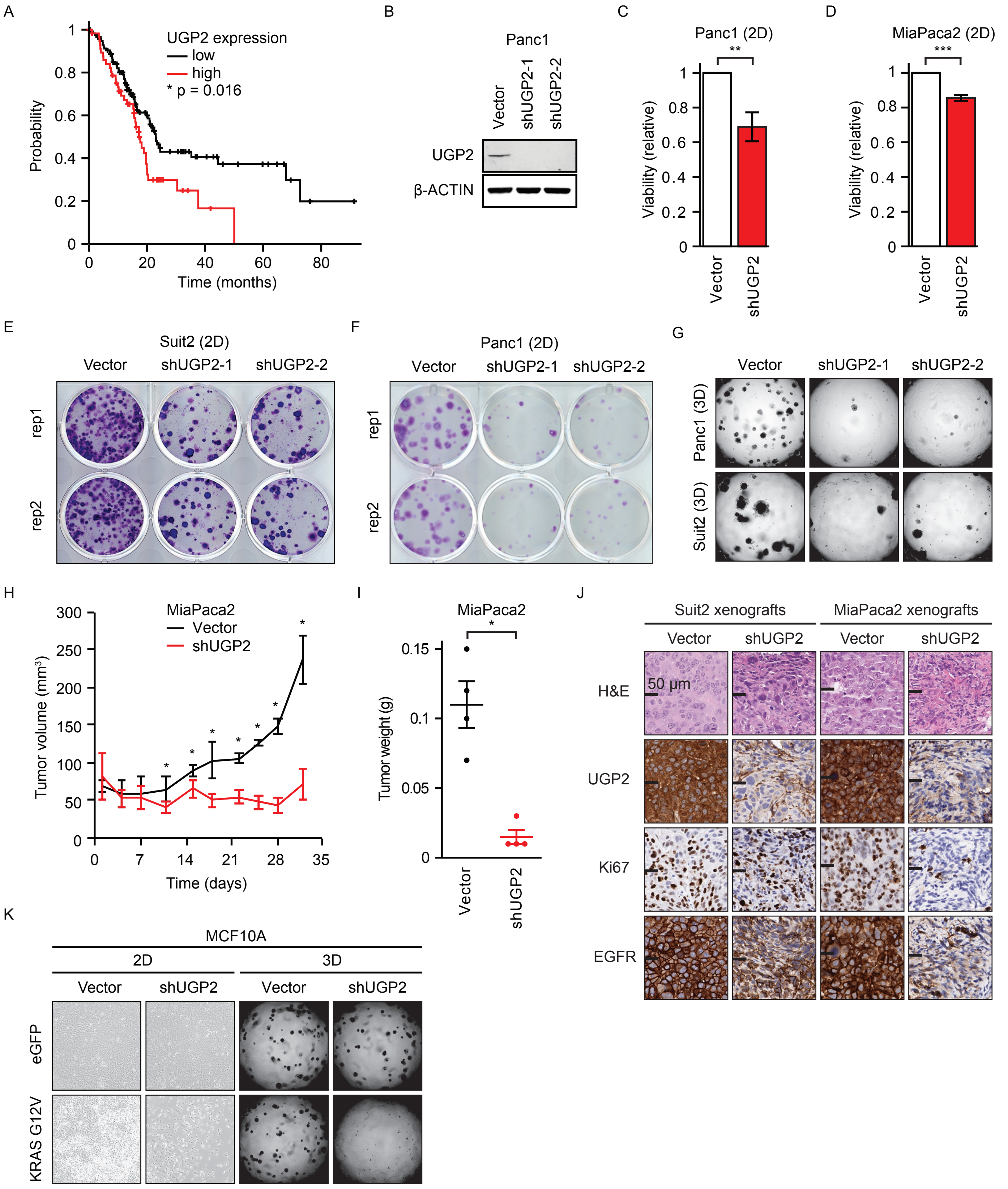


**S1. UGP2 is critical for cell growth in cells with mutant Ras.**

A, Cumulative probability of survival of 177 patients with pancreatic ductal adenocarcinoma split by expression of UGP2 ($p = 0.016$). Data adapted from cBioPortal (Cerami et al., 2012; Gao et al., 2013). B, Immunoblots for shRNAs knocking down UGP2 in Panc1 cells, probed as indicated. C-D, Relative viability by CellTiter-Glo after 48 hours of growth in 2D culture of MiaPaca2 (C) and Suit2 (D) cells with shUGP2 or empty vector control. **$p<0.01$, ***$p<0.001$. E-F, Representative crystal violet staining of Suit2 (E) and Panc1 (F) cells with two independent shRNAs against UGP2 or empty vector control grown in two-dimensional culture for 10 days. G, Representative images of Panc1 and Suit2 cells stably expressing shUGP2 or Vector control grown in three-dimensional matrigel for 14 days. H, Tumor volumes of MiaPaca2 cells with shUGP2 or empty vector control xenografted on opposite flanks of nude mice, n = 4, * $p < 0.05$. I, Tumor weights at endpoint, n = 4, * $p < 0.05$. J, Representative immunohistochemical staining of paired opposite-flank tumors with shUGP2 or vector control at endpoint, stained as indicated. Scale bar represents 50 μM. K, Representative images of MCF10A cells stably expressing shUGP2 or empty vector grown in 2D or 3D matrigel for 14 days, n = 3.

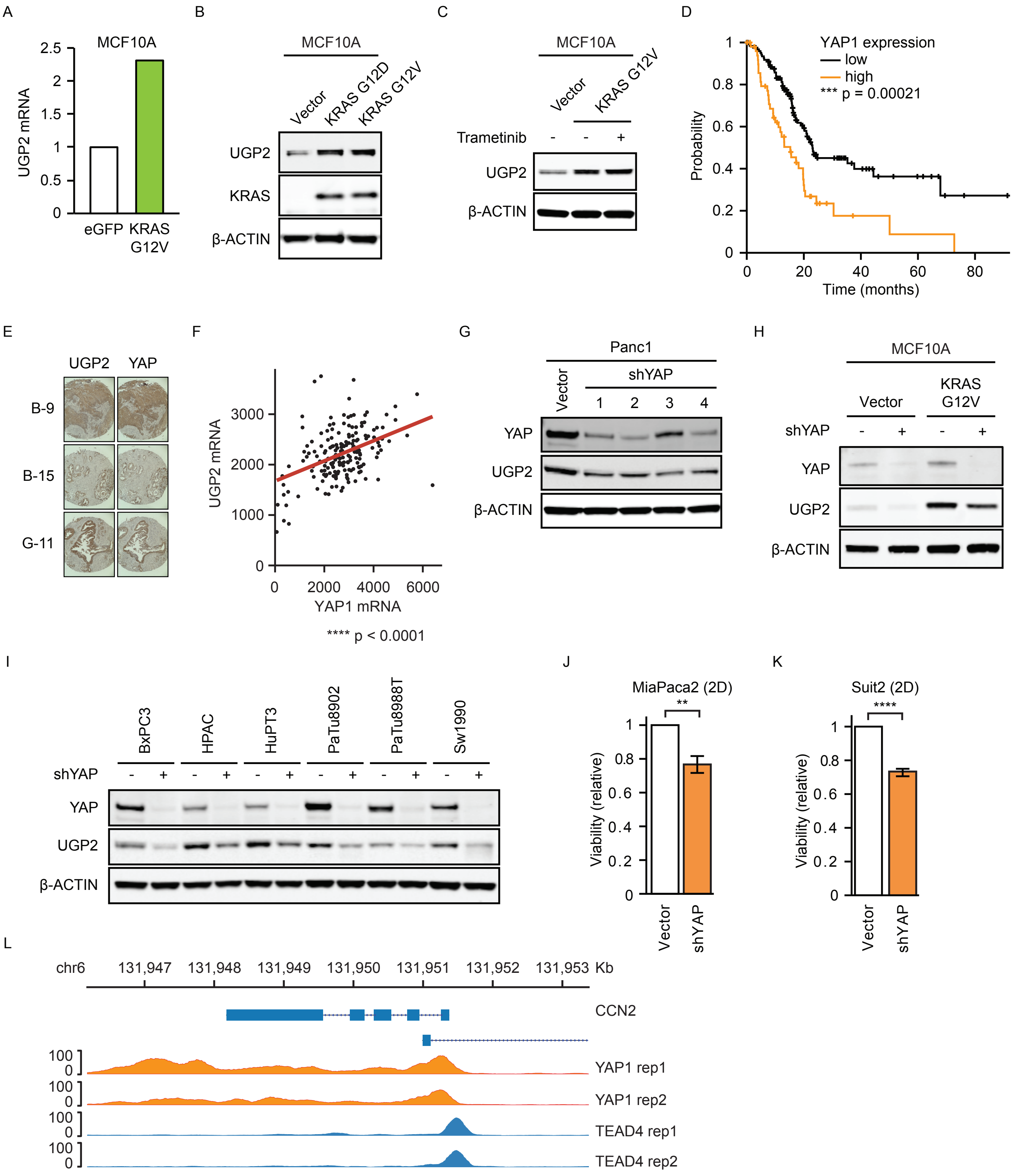


**S2. Regulation of UGP2 by YAP1.**

A, mRNA expression of UGP2 from RNAseq on MCF10A cells stably expressing KRAS G12V or empty vector. B-C, Immunoblots on lysates from MCF10A cells stably expressing KRAS G12D, KRAS G12V, or empty vector (B) or treated with 100 nM trametinib for 48 hours (C). D, Cumulative probability of survival in PDAC patient samples split by expression of YAP1, n = 177, *** p = 0.00021. E, Representative tissue microarray images stained for YAP or UGP2. B-9, B-15, and G-11 indicate sample codes. F, Correlation between UGP2 and YAP1 mRNA in pancreatic ductal adenocarcinoma patient samples. Analysis performed using cBioPortal, n = 178, $R^2$ = 0.18, Pearson = 0.43, p = 3*10-9 (Cerami et al., 2012; Gao et al., 2013). G-I, Immunoblots on cells with shRNAs against YAP or vector control, probed as indicated. Shown are Panc1 with four different YAP shRNAs (G), MCF10A stably expressing KRAS G12V or control eGFP (H), and a panel of pancreatic cancer cell lines (I). J-K, Relative viability of MiaPaca2 (J) and Suit2 (K) cells by CellTiter-Glo after 48 hours of growth in two-dimensional culture in with shYAP or empty vector control. L, ChIP-seq for the control YAP1/TEAD4 binding site in the CCN2 genomic region.

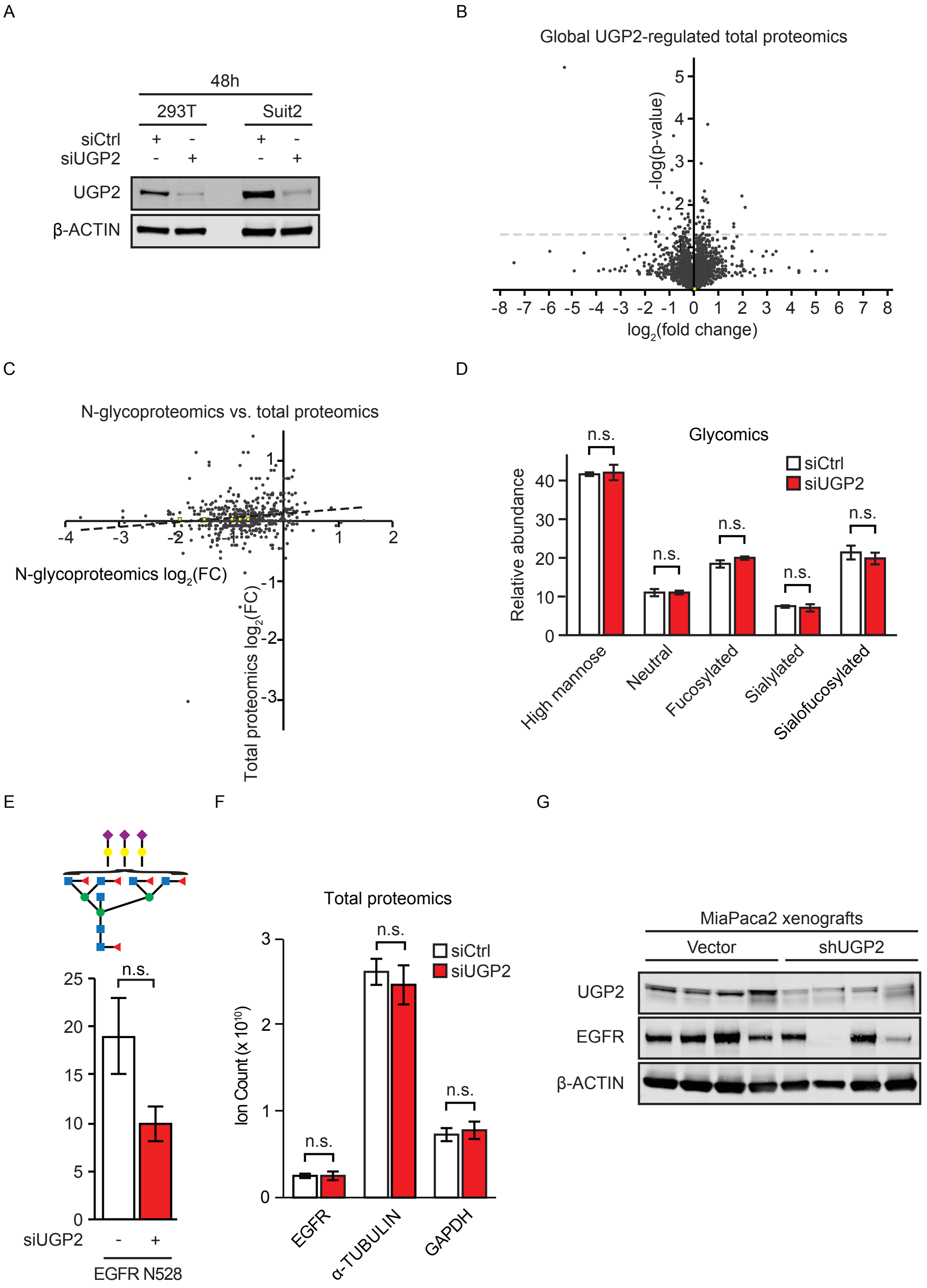


**S3. Glycan changes induced by UGP2 knockdown.**

A, Immunoblots on lysates from 293T or Suit2 cells transfected with siUGP2 or non-targeting control siRNAs for 48 hours, probed as indicated. B, Global total proteomic analysis in Suit2 cells upon knockdown of UGP2 relative to non-targeting control siRNAs after 48 hours, yellow boxes represent total EGFR. C, Shown are $\log_2$ of the fold change in specific modifications plotted against $\log_2$ of the fold change in total protein upon knockdown of UGP2 in Suit2. $R^2 = 0.0215$, yellow boxes represent EGFR modifications. D, Global glycomic comparison in Suit2 cells with siUGP2 of non-targeting control siRNAs at 48 hours, n.s. not significant at a threshold of $p < 0.05$. E, Quantification of glycan modifications on EGFR N528 in Suit2 cells with siUGP2 or non-targeting control siRNAs at 48 hours. n.s. not significant at a threshold of $p < 0.05$, blue squares represent N-acetylglucosamines, green circles represent mannoses, yellow circles represent galactoses, red triangles represent fucoses, and purple diamonds represent N-acetylneuraminic acid. F, Quantification of total EGFR, α-TUBULIN, and GAPDH proteins by mass spectrometry in Suit2 cells upon knockdown of UGP2 or non-targeting control siRNAs. n.s. not significant at a threshold of $p < 0.05$. G, Immunoblot of MiaPaca2 xenograft endpoint tumor lysates, probed as indicated.

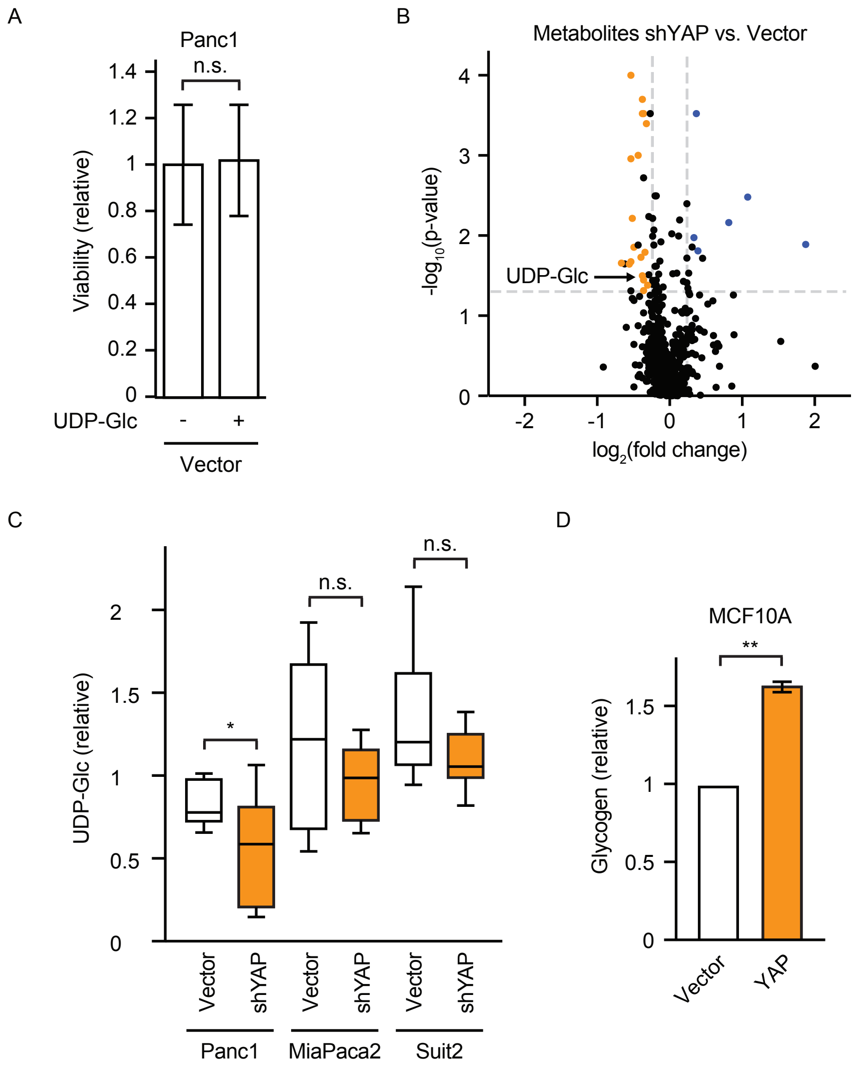


**S4. Functional effects of UGP2 and YAP perturbation.**

A, Relative viability of Panc1 cells with vector control grown with or without 100 μM UDP-Glc for 7 days, n.s. not significant at a threshold of $p < 0.05$. B, Global metabolomics of Panc1 cells with shYAP or empty vector control. Orange dots indicate metabolites in shYAP cells significantly decreased $p < 0.05$, blue dots indicate metabolites significantly increased $p < 0.05$, arrow indicates UDP-Glucose. C, Relative glycogen in MCF10A cells expressing YAP or empty vector control. ** $p < 0.01$. D, UDP-glucose levels or Panc1, MiaPaca2, and Suit2 cells with shYAP or empty vector. * $p < 0.05$ by ANOVA, n.s. not significant.
